# A pan-cancer PDX histology image repository with genomic and pathological annotations for deep learning analysis

**DOI:** 10.1101/2022.10.26.512745

**Authors:** Brian S White, Xing Yi Woo, Soner Koc, Todd Sheridan, Steven B Neuhauser, Shidan Wang, Yvonne A Evrard, John David Landua, R Jay Mashl, Sherri R Davies, Bingliang Fang, Maria Gabriela Raso, Kurt W Evans, Matthew H Bailey, Yeqing Chen, Min Xiao, Jill Rubinstein, Ali Foroughi pour, Lacey Elizabeth Dobrolecki, Maihi Fujita, Junya Fujimoto, Guanghua Xiao, Ryan C Fields, Jacqueline L Mudd, Xiaowei Xu, Melinda G Hollingshead, Shahanawaz Jiwani, PDXNet consortium, Brandi Davis-Dusenbery, Tiffany A Wallace, Jeffrey A Moscow, James H Doroshow, Nicholas Mitsiades, Salma Kaochar, Chong-xian Pan, Moon S Chen, Luis G Carvajal-Carmona, Alana L Welm, Bryan E Welm, Ramaswamy Govindan, Shunqiang Li, Michael A Davies, Jack A Roth, Funda Meric-Bernstam, Yang Xie, Meenhard Herlyn, Li Ding, Michael T Lewis, Carol J Bult, Dennis A Dean, Jeffrey H Chuang

## Abstract

Patient-derived xenografts (PDXs) model human intra-tumoral heterogeneity in the context of the intact tissue of immunocompromised mice. Histological imaging via hematoxylin and eosin (H&E) staining is performed on PDX samples for routine assessment and, in principle, captures the complex interplay between tumor and stromal cells. Deep learning (DL)-based analysis of large *human* H&E image repositories has extracted inter-cellular and morphological signals correlated with disease phenotype and therapeutic response. Here, we present an extensive, pan-cancer repository of nearly 1,000 *PDX* and paired human progenitor H&E images. These images, curated from the PDXNet consortium, are associated with genomic and transcriptomic data, clinical metadata, pathological assessment of cell composition, and, in several cases, detailed pathological annotation of tumor, stroma, and necrotic regions. We demonstrate that DL can be applied to these images to classify tumor regions and to predict xenograft-transplant lymphoproliferative disorder, the unintended outgrowth of human lymphocytes at the transplantation site. This repository enables PDX-specific, investigations of cancer biology through histopathological analysis and contributes important model system data that expand on existing human histology repositories. We expect the PDXNet Image Repository to be valuable for controlled digital pathology analysis, both for the evaluation of technical issues such as stain normalization and for development of novel computational methods based on spatial behaviors within cancer tissues.

## Introduction

The high clinical failure rate of cancer therapies is often attributed to the lack of tumor heterogeneity in pre-clinical models (Tentler et al. 2012). This concern has motivated increased use of patient-derived xenografts (PDXs), in which a fresh human tumor biopsy is implanted subcutaneously or orthotopically in the flank of an immunodeficient mouse. If the implantation successfully establishes the model in the *P*_*0*_ mouse, the tumor can be passaged in future generations (*P*_*1*_, *P*_*2*_, etc.). After being sufficiently expanded, the model may be used for pre-clinical drug trials and these have successfully predicted therapeutic outcome in patients (Pompili et al. 2016).

PDXs have been shown to recapitulate phenotypes of their human progenitors along other dimensions as well. For example, PDX expression profiles correlate with those of their progenitors (Petrillo et al. 2012; S. Li et al. 2013) and these can be stably maintained for at least five generations (X. Zhang et al. 2013). Similar consistency between PDX and human and across passages has been demonstrated for copy number alterations (Woo et al. 2021). Finally, PDXs retain the invasive histological phenotype of their matched progenitor, as reflected in staining with hematoxylin and eosin [H&E; (X. Zhang et al. 2013)].

This latter finding raises the intriguing possibility that recent successes applying deep learning (DL) for whole-slide (WSI) analysis of H&E images in *human* (Aeffner et al. 2019; Komura and Ishikawa 2018; Srinidhi, Ciga, and Martel 2021) will be applicable to PDXs, as well. DL-based analyses of human data are capable of predicting metastases (Ehteshami Bejnordi et al. 2017), gene mutations (Coudray et al. 2018) and expression (Coudray et al. 2018; Schmauch et al. 2020), survival (Kather et al. 2019), cancer types (Noorbakhsh et al. 2020), molecular (Sirinukunwattana et al. 2021), clinical (Rawat et al. 2020), and histological (J. Li et al. 2022) tumor subtypes, and response to both chemotherapy (Farahmand et al. 2022) and immune checkpoint inhibitors (Johannet et al. 2021), amongst many other phenotypes reviewed extensively elsewhere (Tran et al. 2021; van der Laak, Litjens, and Ciompi 2021; Bera et al. 2019). DL approaches can, in principle, assess the complexity of spatial interactions between cancer, stromal, and immune cells reflected in H&E images, thus moving beyond cancer cell-intrinsic, univariate gene biomarkers. For example, DL methods were able to associate tumor-infiltrating lymphocyte spatial structure with survival (Saltz et al. 2018), and differentiate breast tumor samples from benign breast biopsies based on stroma (Saltz et al. 2018; Ehteshami Bejnordi et al. 2018).

The explosive growth of DL-based applications in digital pathology has been made possible by extensive, public repositories of human H&E images, such as in TCGA, The Cancer Imaging Archive, and Imaging Data Commons. No similar resource exists for PDX images. Here, we describe a large-scale repository of >800 PDX and >100 matched patient tumor H&E images, along with expansive genomic, transcriptomic, clinical, and pathological annotations. The images were curated as part of the National Cancer Institute’s PDX Development and Trial Centers Research Network program (PDXNet), aimed at collaborative development and pre-clinical testing of targeted therapeutic agents. Thumbnails and clinical metadata of the images, along with the raw genomic and transcriptomic data, are avaliable on the PDXNet portal (Koc et al. 2022). The raw images are hosted on the Seven Bridges Cancer Genomics Cloud [CGC; www.cancergenomicscloud.org; (Lau et al. 2017)] and will soon be available on the Imaging Data Commons (Fedorov et al. 2021). We present several use cases applying DL for classification of PDX images. The repository should facilitate monitoring of potential divergence between a PDX and its human progenitor during passaging (Ben-David et al. 2017) and exploration of questions uniquely relevant to PDX models, including the impact of human cell turnover within the xenografts (Blomme et al. 2018; Sanz et al. 2009; Gray et al. 2004). We expect this PDXNet Image Repository to be valuable for controlled digital pathology analysis and development of novel computational methods based on spatial behaviors within patient-derived cancer tissues.

## Results

### A pan-cancer, multi-institutional repository links histology images to clinical annotations, pathological assessments, and genomic and transcriptomic data

The PDXNet repository consists of 810 H&E images from PDXs and 139 from their paired human progenitors. They represent 33 cancer types (**Fig. 1A**), including breast cancer (BC; a category encompassing ductal carcinoma *in situ*, lobular carcinoma *in situ*, invasive breast carcinoma, invasive lobular carcinoma, and breast cancer not otherwise specified; *n*=134), colon adenocarcinoma (COAD; *n*=94), pancreatic cancer (PDAC; *n*=87), lung adenocarcinoma (LUAD; *n*=80), melanoma (SKCM; *n*=71), and squamous cell lung cancer (LUSC; *n*=67). Images were generated at five PDX Develoment and Trial Centers (PDTCs) within PDXNet – Baylor College of Medicine (BCM), Huntsman Cancer Institute (HCI), MD Anderson Cancer Center (MDACC), The Wistar Institute (WISTAR), and Washington University in St Louis (WUSTL) – as well as at the NCI Patient-Derived Models Repository (PDMR) and The Jackson Laboratory (JAX).

**Fig 1.**
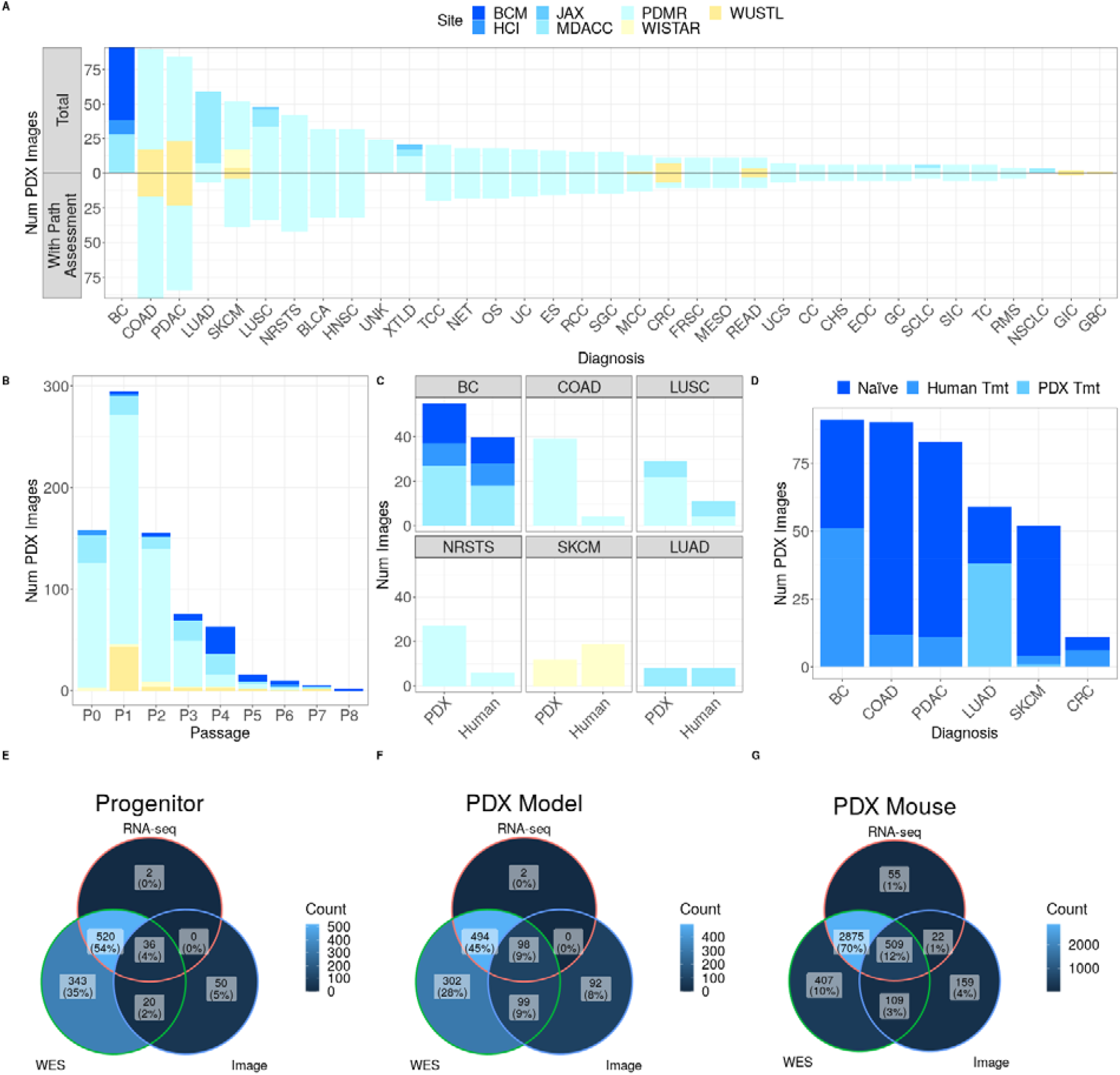
PDXNet Image Repository captures heterogeneity across tumor types, passages, and treatment status and pairs histology images with pathological assessment and genomic and transcriptomic data. (A) Distribution of total PDX H&E images and subset with pathological assessment by diagnosis. (B) Distribution of PDX images by passage. (C) Counts of images by source for those derived from human progenitor / PDX pairs, restricted to diagnoses with at least four such progenitors. (B, C) Barcharts colored according to contributing site as indicated in (A). (D) Distribution of PDX images across diagnoses with at least two treated PDXs or PDXs derived from at least two treated progenitors. (E, F, G) Intersection of image, RNA-seq, and WES data (E) amongst human progenitor samples, (F) across PDX models, and (G) across individual mice.

Most PDX images are from early passages, with those from P0 (*n*=158), P1 (*n*=295), and P2 (*n*=155) predominating over those from later passages (> P2; *n*=171; **Fig. 1B**). 102 images from human progenitors have at least one paired PDX image, for a total of 306 such images. Of these, cancer diagnoses with images from at least four unique, paired progenitors include BC, COAD, LUSC, Non-Rhabdomyosarcoma soft tissue sarcomas (NRSTS; a category encompassing non-infantile fibrosarcoma, gastrointestinal stromal tumor, non-uterine leiomyosarcoma, malignant peripheral nerve sheath tumor, and synovial sarcoma), SKCM, and LUAD (**Fig. 1C**). The repository includes images from mice that were treated therapeutically, whose human progenitor was treated therapeutically, or that are completely treatment naïve. Diagnoses having at least two treated PDXs or PDXs derived from at least two treated progenitors include BC, COAD, PDAC, LUAD, SKCM, and colorectal cancer (CRC; **Fig. 1D**).

H&E images are linked to RNA-seq and whole exome sequencing (WES) data hosted on the PDXNet portal (Koc et al. 2022): 36 of 106 human progenitors, 98 of 289 PDX models, and 509 of 799 mice bearing PDXs are characterized by histological imaging, transcriptomically, and genomically (**Figs 1E-G**). Additionally, most images include pathological assessment of tumor stage (*n*=650) and slide-level proportions of cancer, stromal, and necrotic regions (*n*=626; **Fig. 1A**).

### Pathologist cell type estimates allow evaluation of cell segmentation and classification

We used pathologist estimates of percentage of neoplastic, stromal, and necrotic cells in *PDX* samples as ground truth in evaluating Hover-Net, a nuclei segmentation and classification trained on *human* H&E images (Graham et al. 2019). Hover-Net classifies nuclei as neoplastic, stromal (“connective”), necrotic, immune (“inflammatory”), non-neoplastic epithelial, and other. We derived a slide-level estimate for each cell type as the proportion of the corresponding nuclei area relative to the total tissue area (see Methods). Hover-Net-based slide-level estimates generally correlated well for neoplastic (median *weighted* Pearson correlation *r* = 0.51), stromal (*r* = 0.52), and necrotic (*r* = 0.70) cells across diagnoses (COAD, CRC, and PDAC) and datasets (PDMR and WUSTL; **Fig. 2**).

**Fig 2.**
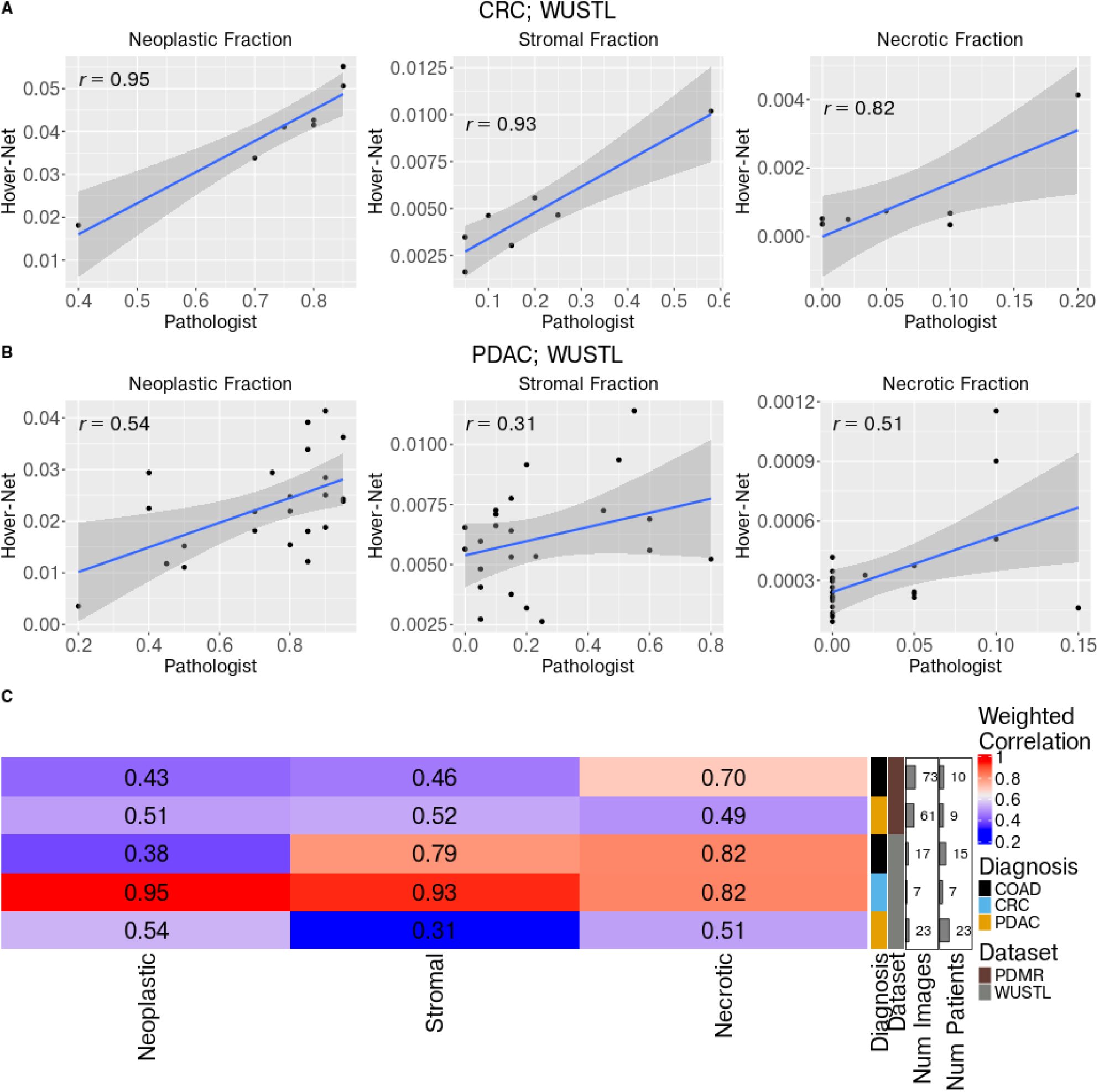
Cell segmentation and classification trained on human data correlate with pathologist estimates of cell type proportions in PDX samples. (A, B) Correlation of pathologist (x axis) and Hover-Net-based slide-level estimates (y) axis for neoplastic (left column), stromal (center column), and necrotic (right column) cell proportions in (A) CRC and (B) PDAC from WUSTL. Each point corresponds to an individual image and PDX (i.e., no replication). (C) Weighted Pearson correlations of pathologist and Hover-Net-based estimates of neoplastic, stromal, and necrotic cell proportions in COAD, CRC, and PDAC in the PDMR and WUSTL datasets. Correlations are computed over images, inversely weighted according to number of replicate images derived from the same patient. In cases without replication, the number of images equals the number of patients and weighted and unweighted Pearson correlation are identical.

### Pathologist pixel-level annotations enable training of a regional classifier

To spatially contextualize slide-level correlations between predicted and assessed cell type proportions, a board-certified pathologist manually annotated cancer epithelium (“tumor”), stroma, and necrotic regions within H&E images of LUSC (*n*=15), LUAD (*n*=18), and BC (*n*=8; **Fig. 3E**). We labeled tiles according to their intersection with these annotations and found that, as expected, tumor tiles were predominantly composed of cells classified as neoplastic in the lung datasets (**Fig. S1**). Consistently, the spatial organization of predicted tumor cells closely resembled pathologist annotations (**Fig. 3A**). Breast cancer exhibited a different pattern, with most tiles composed of a mix of neoplastic and stromal cells. Lung stromal tiles harbored predicted neoplastic, immune, and stromal cells, whereas breast stromal tiles were predominantly comprised of stromal cells. Finally, necrotic tiles had levels of predicted necrotic cells in excess of those of other tiles in both tumor types, despite the additional presence of a larger proportion of neoplastic and immune cells in lung cancer and an appreciable proportion of stromal cells in breast cancer. To begin to address cellular heterogeneity within the stromal and necrotic regions, we applied a second nuclei segmentation and phenotyping approach, HD-Staining (S. Wang et al. 2020), to a PDX lung sample (**Fig. 3B**). The spatial pattern of HD-Staining phenotypes was broadly similar to that predicted by Hover-Net – though HD-Staining’s prediction of macrophages within one necrotic area was more fine-grained than the immune prediction of Hover-Net and the two disagreed about the immune content with one stromal region.

**Fig 3.**
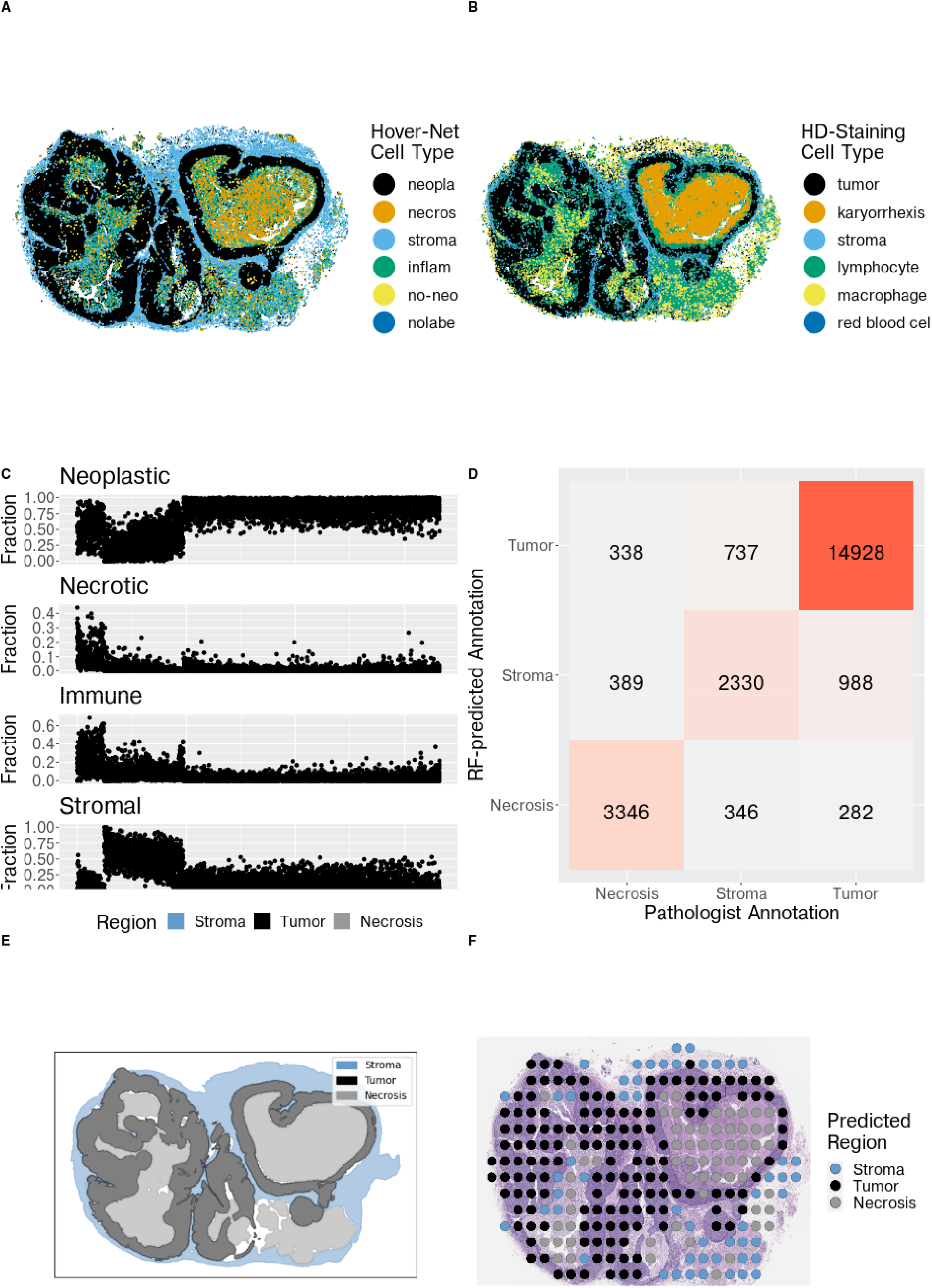
Regional classifier predicts cancer epithelium (“tumor”), stroma, and necrotic areas based on nuclei classification. (A, B) Cell phenotypes predicted by (A) Hover-Net or (B) HD-Staining within a PDX lung tumor. (C) Labeled tiles (columns; blue: stroma, black: tumor, gray: necrosis) represented according to the cellular fraction that is predicted by Hover-Net to be neoplastic, necrotic, immune, or stromal. (D) Confusion matrix comparing pathologist-provided tile labels with corresponding prediction from the RF Hover-Net classifier. (E) Pathologist regional annotations. (F) Random forest (RF) Hover-Net-based classifier predictions of regions (within 512 × 512 pixels).

**Fig S1.**
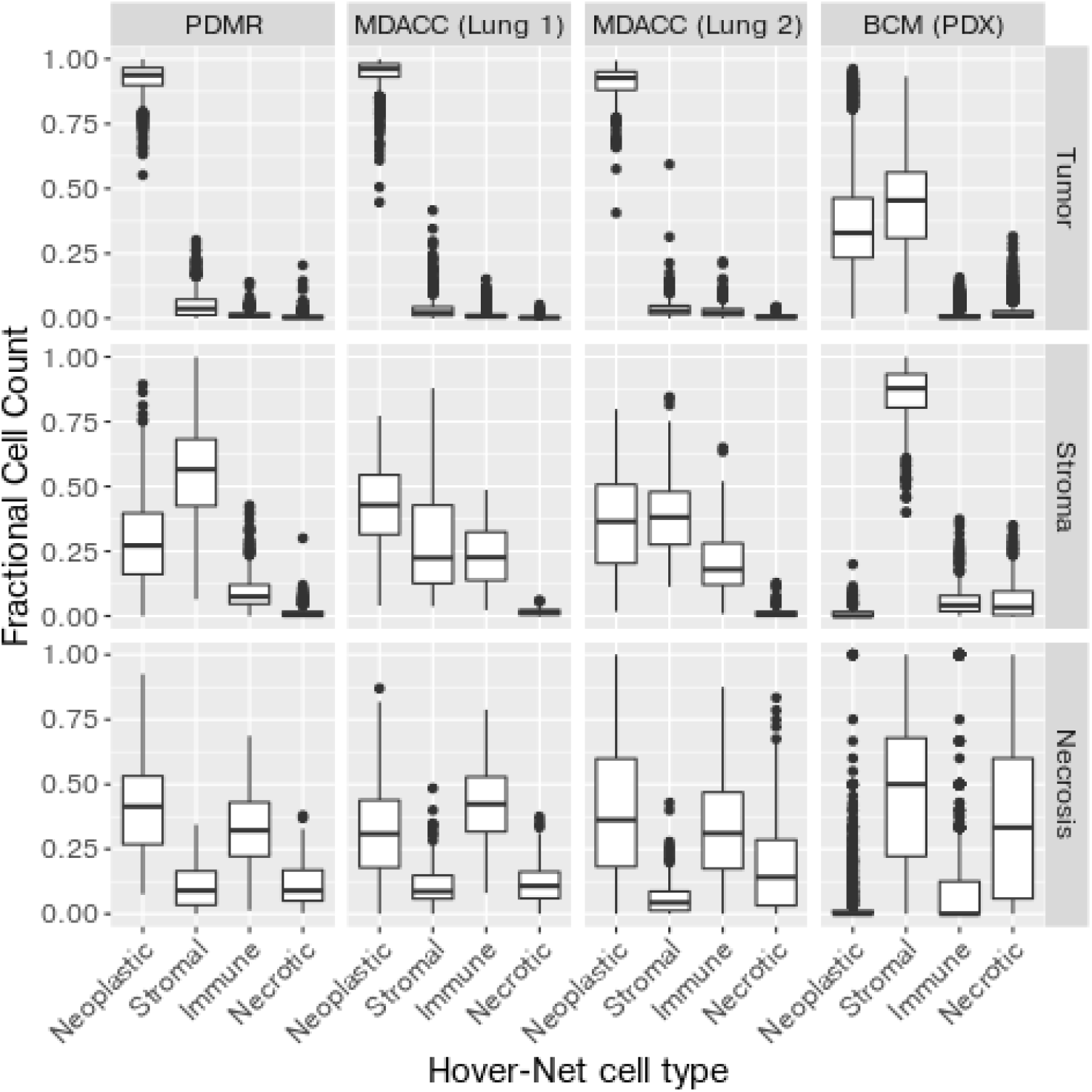
Regional proportions of cell types vary by cancer type. Boxplots show fraction of cells (y axis) of a particular type predicted by Hover-Net (x axis) relative to total number of segmented cells within a 512×512 pixel tile at 20X resolution. Results are stratified by dataset (columns) and pathologist-annotated region overlapping the tile (rows).

The localization of particular cell types within morphological regions has clinical implications. For example, tumor-stromal interactions have been associated with disease progression and chemotherapy resistance in breast cancer (Farmer et al. 2009; Conklin and Keely 2012; Criscitiello, Esposito, and Curigliano 2014; “The Impact of Tumor Stroma on Drug Response in Breast Cancer” 2015; Plava et al. 2019). While regional annotations facilitate correlative studies with clinical phenotypes, the manual burden in generating them is high: a pathologist expended approximately 20 hours demarcating, in detail, regions within each sample.

To alleviate this annotation burden, we instead sought to predict tumor, stroma, and necrotic regions within H&E images. We first represented each tile according to the fraction (**Fig. 3C**) and count (**Fig. S2**) of each Hover-Net-predicted cell type within it. Consistent with our previous results, we observed that these tile profiles of constituent cell types were region- and cancer type-specific. We used the profiles as features and trained a random forest to (probabilistically) predict regions using them. Predictions correctly recapitulated irregular region boundaries (**Fig. 3F**) and performance was high (**Fig. 3D**; precision, recall, specificity, and accuracy all equal 0.87). Further, prediction performance was consistently high for all regions across cancer types, datasets, and individual samples: over tiles labeled as belonging to a particular region, the median probability of being assigned to that correct region was over 50% for the four samples in the lung PDMR dataset, for all but two of ten lung stromal regions and one of 12 tumor lung tumor regions in the MDACC dataset, and for all but one of seven breast tumor regions and one of five breast necrosis regions in the BCM dataset (**Fig. S3**).

**Fig S2.**
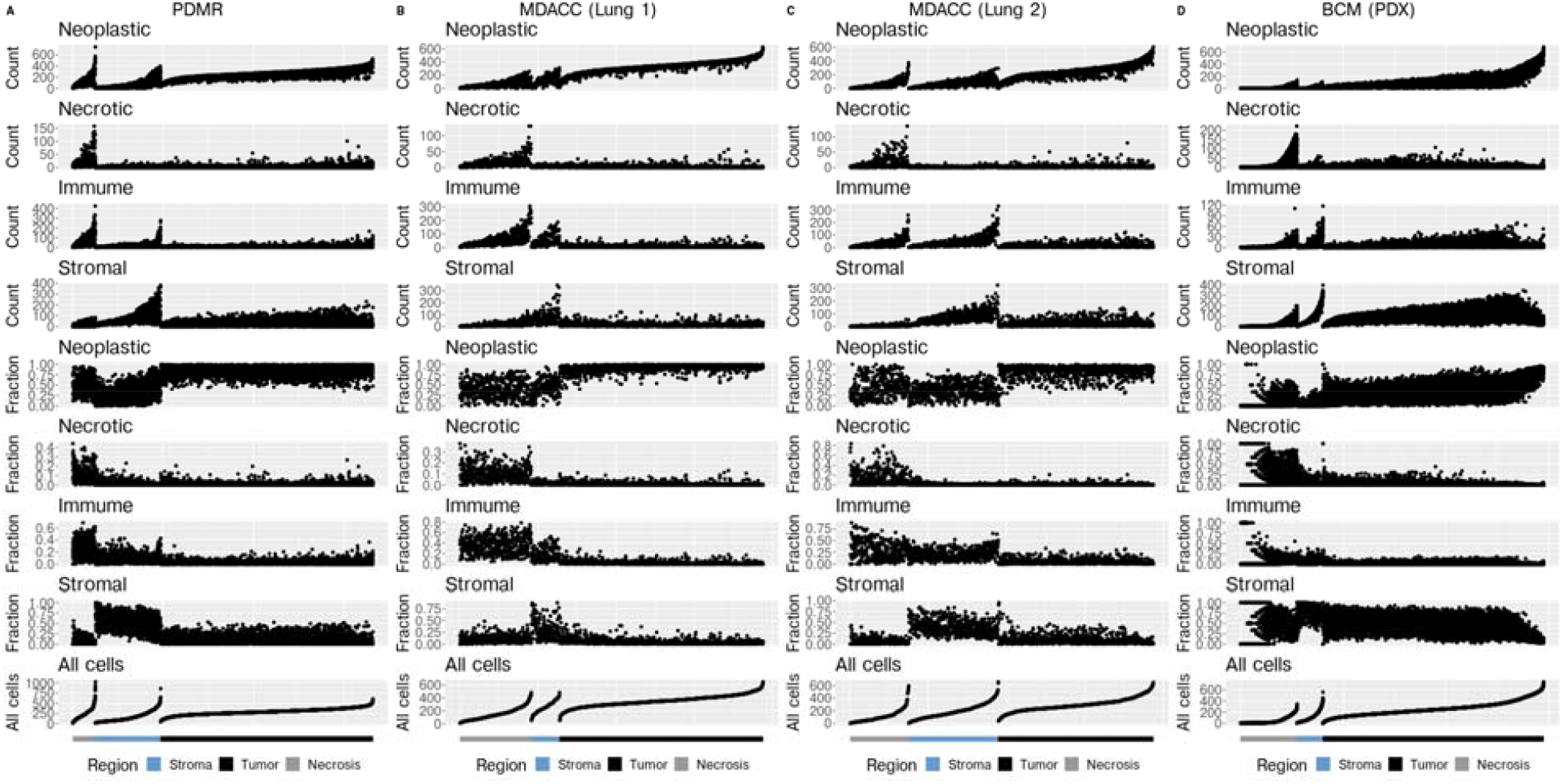
Regions have characteristic cell type proportions and counts that vary by cancer type. Each tile (columns within subpanels) is annotated according to region (tumor, stroma, or necrotic) and represented by the count and fractional proportion of each type of cell (neoplastic, necrotic, immune, or stromal) within it, as well as the total number of cells. Tiles are shown for lung cancer samples in the (A) PDMR and (B, C) two MDACC datasets and for breast cancer samples in the (D) BCM dataset.

**Fig S3.**
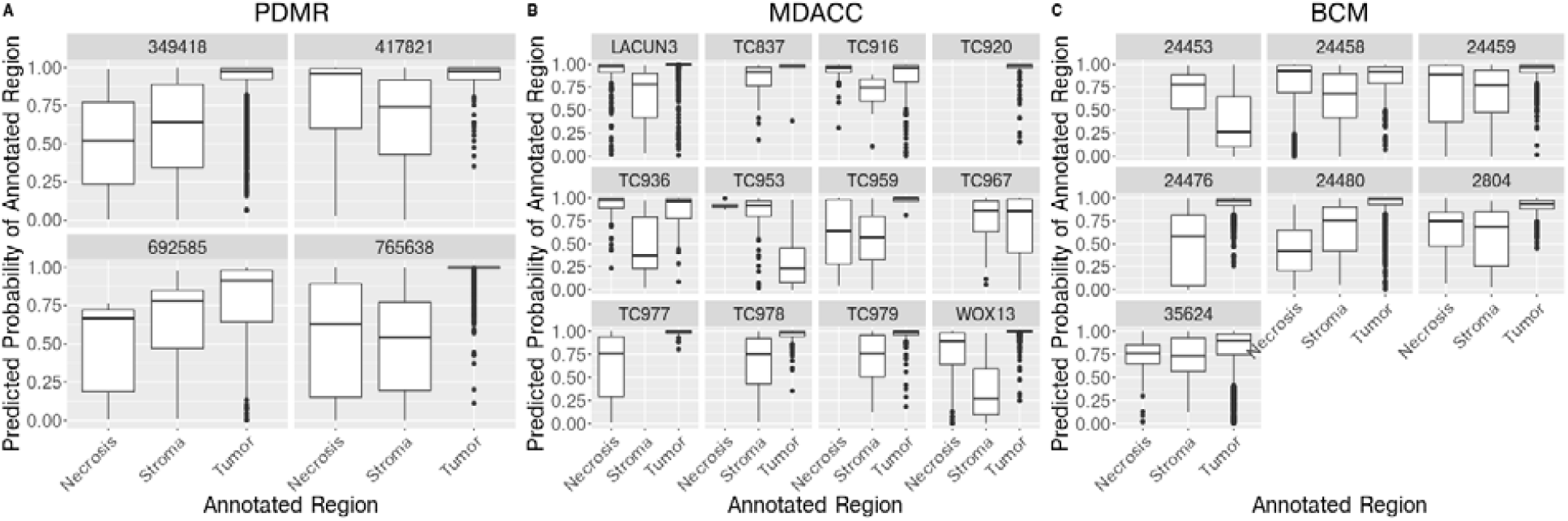
Regional classifier consistently and correctly predicts regions across cancer types, datasets, and samples. Boxplots show distribution over tiles of classifier-predicted probability that tile is assigned to the correct annotated region.

### An H&E-based classifier identifies xenograft-transplant lymphoproliferative disorder

During development of PDX models, human lymphocytes within the implanted tumor biopsy can overtake or limit the growth of the malignant cells. Such xenograft-transplant lymphoproliferative disorder (XTLD) is frequent, occurring at rates that exceed 10% in some transplanted cancer types (Butler et al. 2017; Bondarenko et al. 2015; Dieter et al. 2017; Choi et al. 2016) and that are variable across cancer types (L. Zhang et al. 2015). It can be detected during a quality control (QC) assessment of the generated model using immunohistochemistry (IHC) directed at human pan-immune (CD45) or B-cell (CD20) markers. It has also been detected and predicted based on expression differences between Epstein-Barr virus-associated lymphoma samples and non-lymphoma samples (Woo et al. 2019). Left undetected, however, XTLD would masquerade as the tumor type of the original biopsy, thus invalidating any downstream functional analysis or clinical correlation attempting to characterize that tumor type.

For labs that do not routinely perform IHC during a QC step and for legacy data without such an evaluation, we sought to develop an XTLD classifier applicable to H&E images generated during a standard pathological assessment. We reasoned that the higher density of small lymphocytes characteristic of XTLD (**Fig. 4B**) relative to an epithelial tumor (**Fig. 4A**) would be detectable via cell segmentation and classification. Indeed, we observed that the proportion of segmented immune cells was elevated in XTLD samples relative to LUSC samples in the MDACC (*p* = 4.6 × 10-4) and PDMR (*p* = 0.08) datasets (**Fig. S4A**). Further, median cell area (in 2D sections) of LUSC samples were higher than XTLD samples in MDACC and for all three of five LUSC samples in PDMR (**Fig. S4B**).

**Fig 4.**
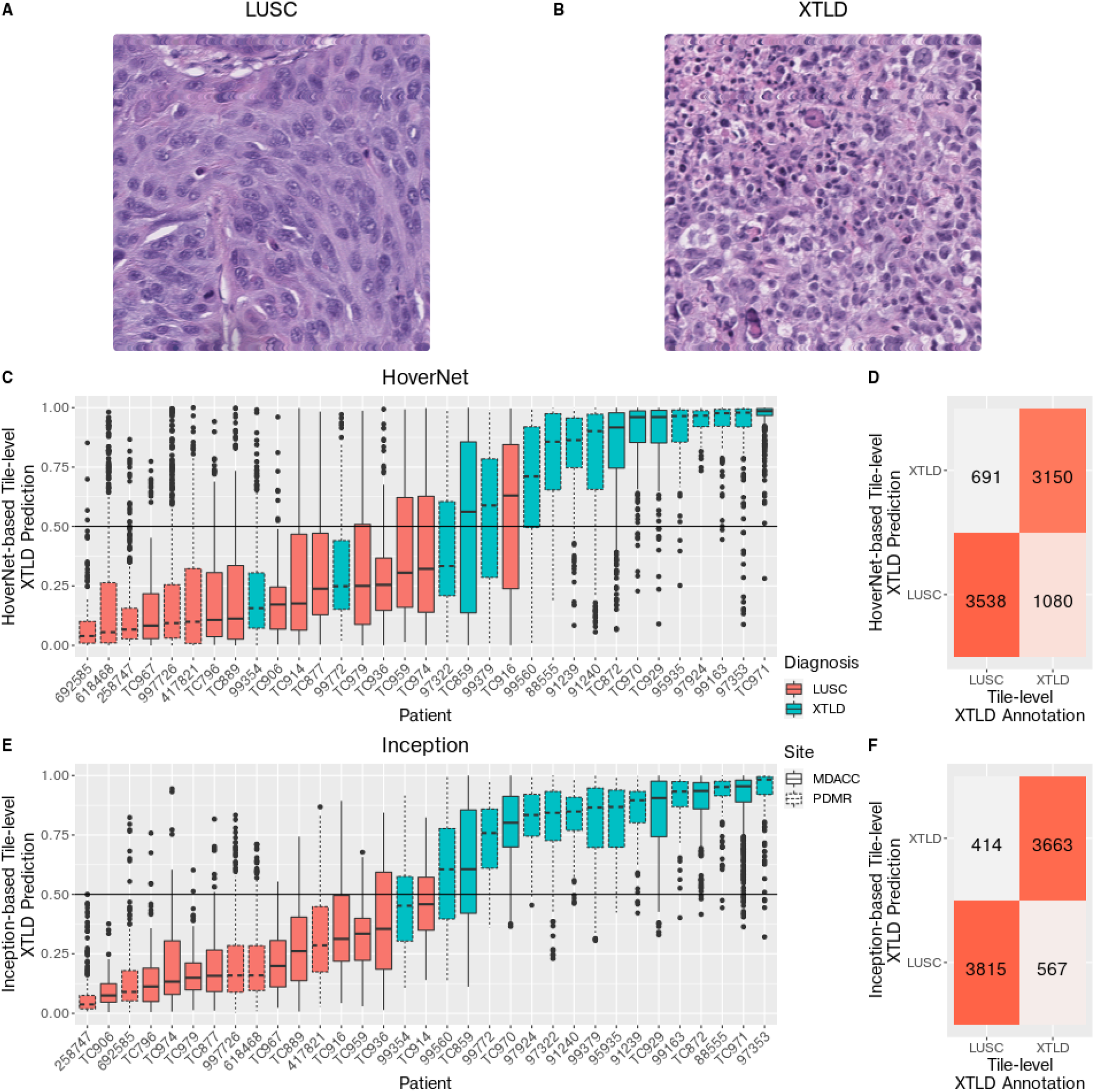
XTLD can be computationally predicted from H&E images using segmented and classified cells or deep learning features. (A, B) H&E images of (A) LUSC and (B) XTLD. (C, D) Hover-Net-based classifier performance. (C) Distribution of tile-level XTLD probability predicted by random forest trained using Hover-Net-derived features, according to patient (x axis) diagnoses with LUSC (red) or XTLD (blue) in the MDACC (solid line) or PDMR (dashed line) datasets. Results shown for held-out set following 5-fold cross validation. (D) Confusion matrix showing tile-level concordance between Hover-Net-based classification (rows) and annotation (columns). (E, F) Inception-based classifier performance, as in (C, D), respectively, but with predictions from a random forest trained using Inception features.

Despite the associations between XTLD and both immune cell proportion and cell size, the orderings induced by these two metrics imperfectly segregate XTLD and LUSC samples. Hence, we attempted a more general classification at the image tile level by representing each tile according to its cell classification-derived proportions of neoplastic, stromal, necrotic, and immune cells, i.e., excluding the count-based features used in the regional classifier. We used these tile-level representations to train a random forest to predict XTLD status, similar to our approach above in predicting region labels. Tile-level prediction performance was strong (**Fig. 4D**), with precision, recall, specificity, accuracy, and AUC between 0.82 and 0.93 (**Fig. S5B, C**). Further, the distribution of tile-level XTLD probabilities were strongly skewed towards one for tiles from XTLD images and towards zero for those from LUSC images (**Fig. S5A**). As expected, immune proportions were more important than neoplastic and necrotic cell proportions and of similar importance to stromal cell proportions in predicting XTLD (**Fig. S6**). At the sample level, the Hover-Net-based classifier predicted the majority of tiles correctly (i.e., with probability > 0.5 for the annotated label) for 29 of 33 samples (**Fig. 4C**).

To improve upon sample level prediction, we next sought to generalize our classifier beyond its four biologically-inspired, but constrained, feature inputs. To do so, we extracted 2048 deep learning (DL) features from each tile using an Inception network v3 pre-trained on ImageNet (Szegedy et al. 2016; Noorbakhsh et al. 2020). These features were then input to a random forest classifier and the classifier was trained using the same assignment of tiles to folds as employed for the Hover-Net-based classifier. The Inception-based classifier improved performance over the Hover-Net-based model at both the tile-level (**Fig. 4F**), with precision, recall, specificity, accuracy, and AUC between 0.94 and 1.00 (**Fig. S5E, F**) and with a more pronounced skew of predicted probabilities towards their correct assignments (**Fig. S5D**), and at the sample-level, where 31 of 33 samples were predicted correctly (**Fig. 4E**). Additionally, the Inception-based model perfectly classified six samples in an external validation dataset (**Fig. 5A**).

**Fig 5.**
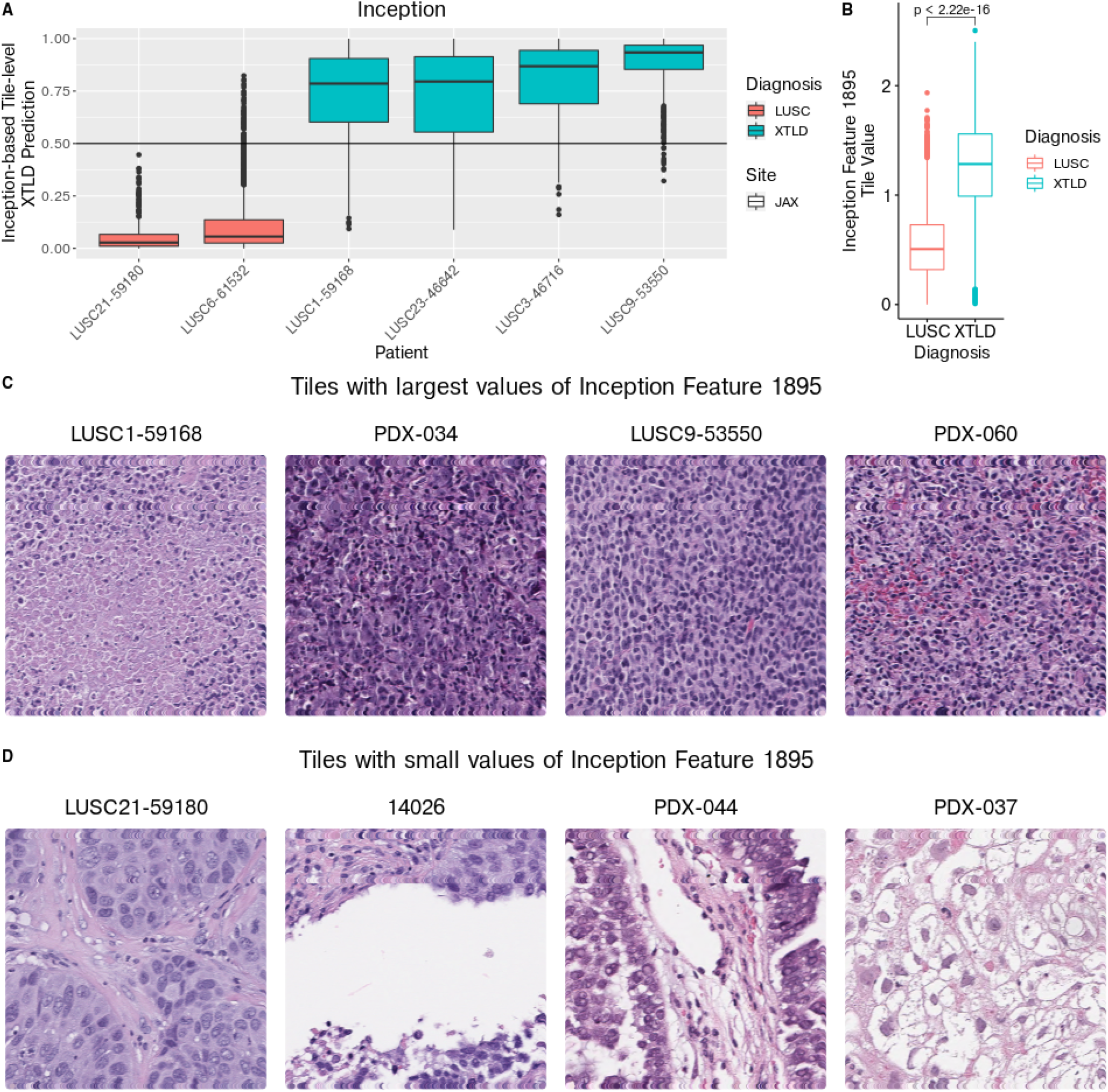
Inception encodes morphological features useful in predicting XTLD. (A) Distribution of tile-level XTLD probability predicted by random forest trained using Inception features, according to patient (x axis) diagnoses with LUSC (red) or XTLD (blue) in The Jackson Laboratory (JAX) external validation dataset. (B) Distribution of most important Inception feature (1895) across tiles within LUSC (red) or XTLD (blue) images. (C) Tiles with large values of Inception feature 1895. The tile with largest value is shown for each of four samples (noted above tile) with highest values, all of which are XTLD cases. (D) Tiles with small values of Inception feature 1895. The tile with smallest value is shown for each of four samples (noted above tile) with smallest values, all of which are LUSC cases.

**Fig S4.**
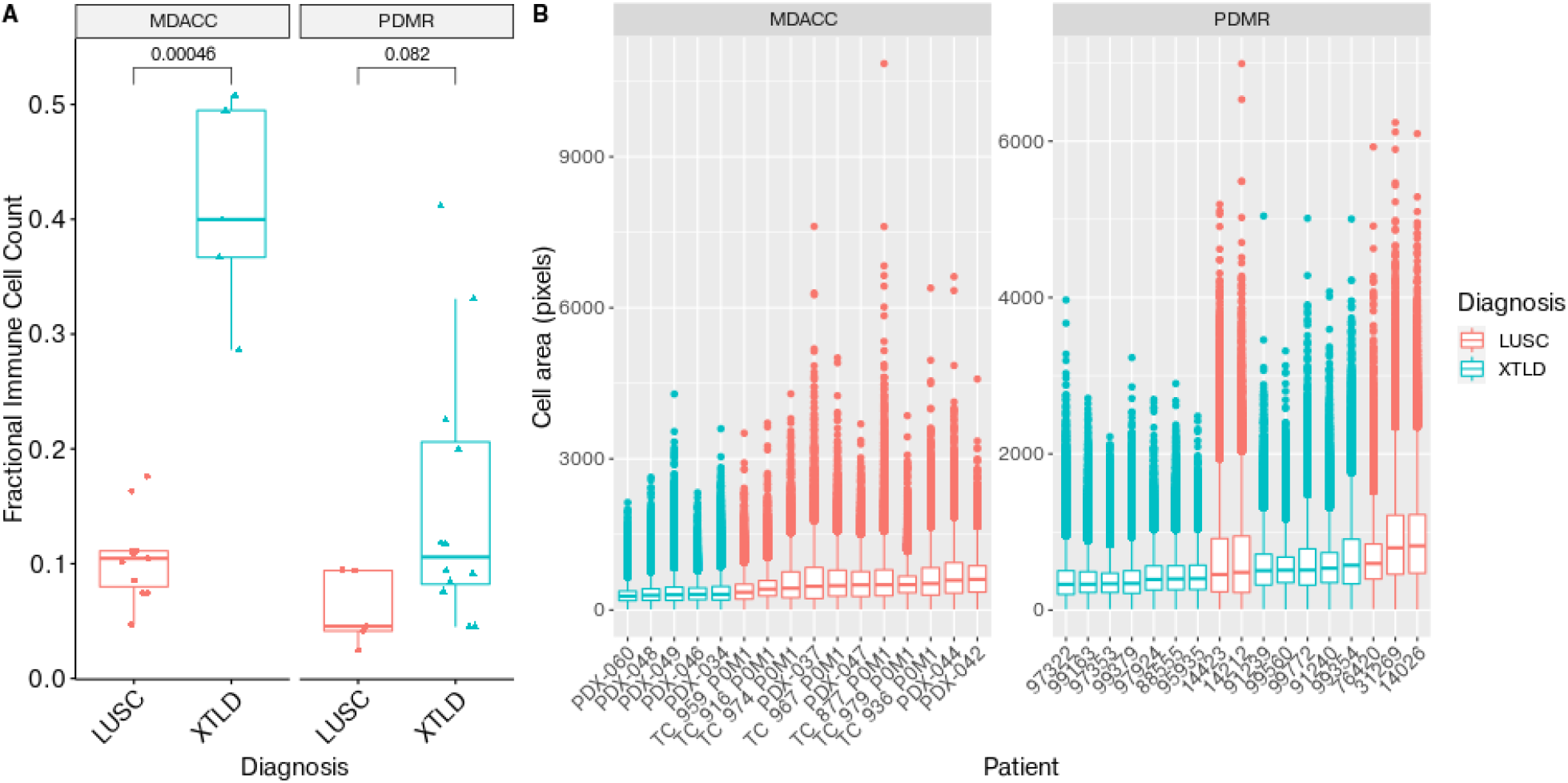
XTLD models have higher immune cell proportion and smaller cell size than LUSC models. (A) Proportion of immune cells relative to all segmented cells (y axis) in LUSC models (red) and XTLD models (blue) within MDACC and PDMR datasets. (B) Distribution of cell area (in pixels; y axis) across LUSC (red) and XTLD (blue) samples (x axis) in MDACC and PDMR datasets.

**Fig S5.**
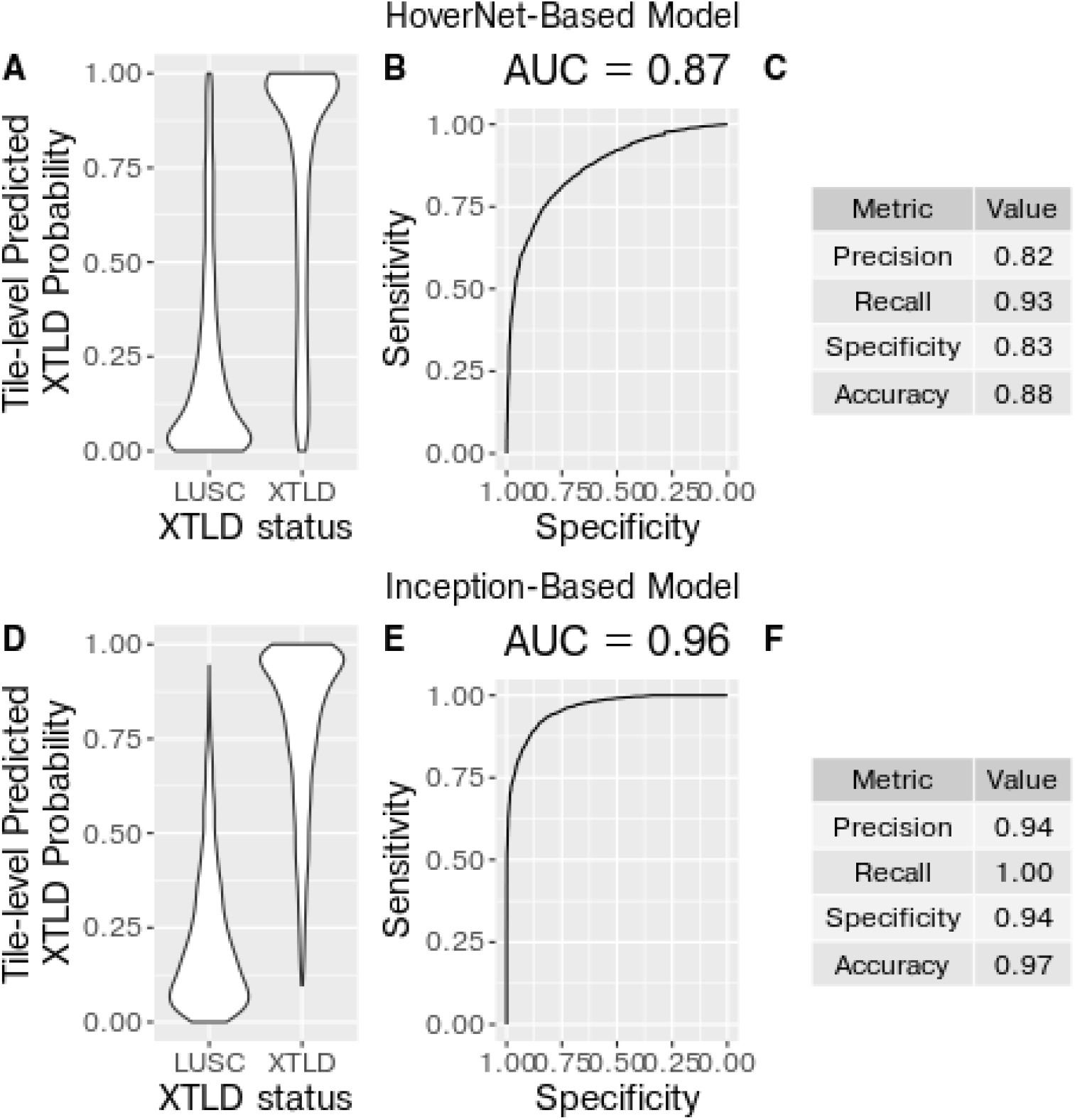
Inception-based classifier outperforms Hover-Net-based classifier in predicting XTLD. Performance of (A, B, C) Hover-Net-based and (D, E, F) Inception-based random forest classifiers of XTLD status. (A, D) Distribution of XTLD probability over tiles in images annotated as LUSC or XTLD as predicted by random forest trained on (A) Hover-Net or (B) Inception features. (B, E) AUC curves over tiles predicted (B) Hover-Net-based or (E) Inception-based classifier. (C, F) Tile-level performance metrics for (C) Hover-Net-based or (F) Inception-based classifier.

**Fig S6.**
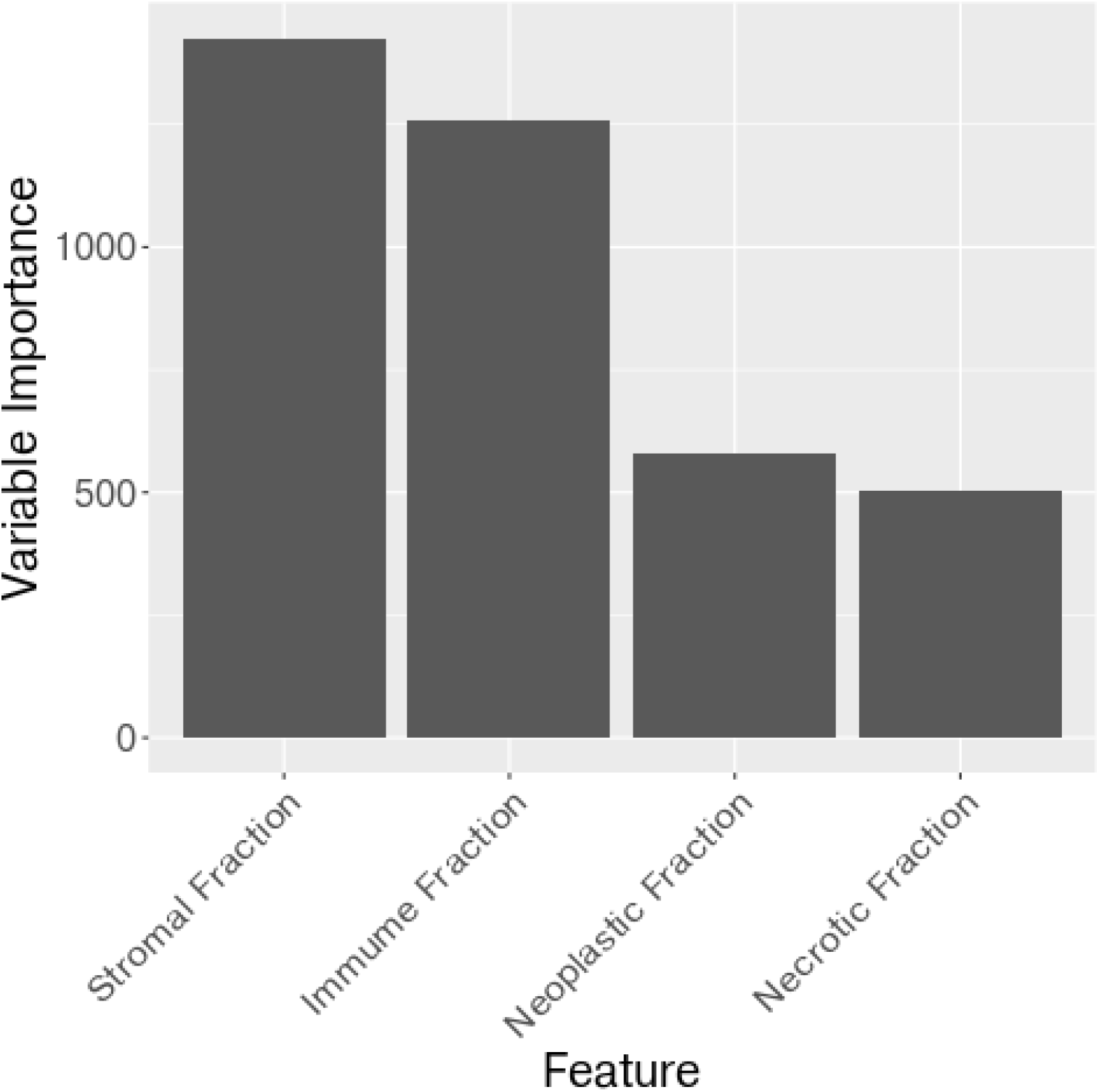
Stromal and immune proportions are more important than neoplastic and necrotic cell proportions in predicting XTLD. Variable importance of Hover-Net-based random forest classifier over tiles represented by proportion of segmented stromal, immune, neoplastic, and necrotic cells.

To interpret the information encoded by Inception that drives classification performance (Foroughi Pour et al. 2022), we examined feature 1895, the feature with highest variable importance in the random forest model (**Fig. S7**). We observed that this single feature segregated tiles in XTLD cases (high feature value) from those in LUSC cases (low feature value; **Fig. 5B**). A montage of tiles suggest that feature 1895 encodes cellular density and/or cell size (or something correlated with these) – those with high feature 1895 value (**Fig. 5C**) are dense in (small, round) lymphocytes or necrotic regions, while those with low value (**Fig. 5D**) have fewer, large cells and admixed stroma.

**Fig S7.**
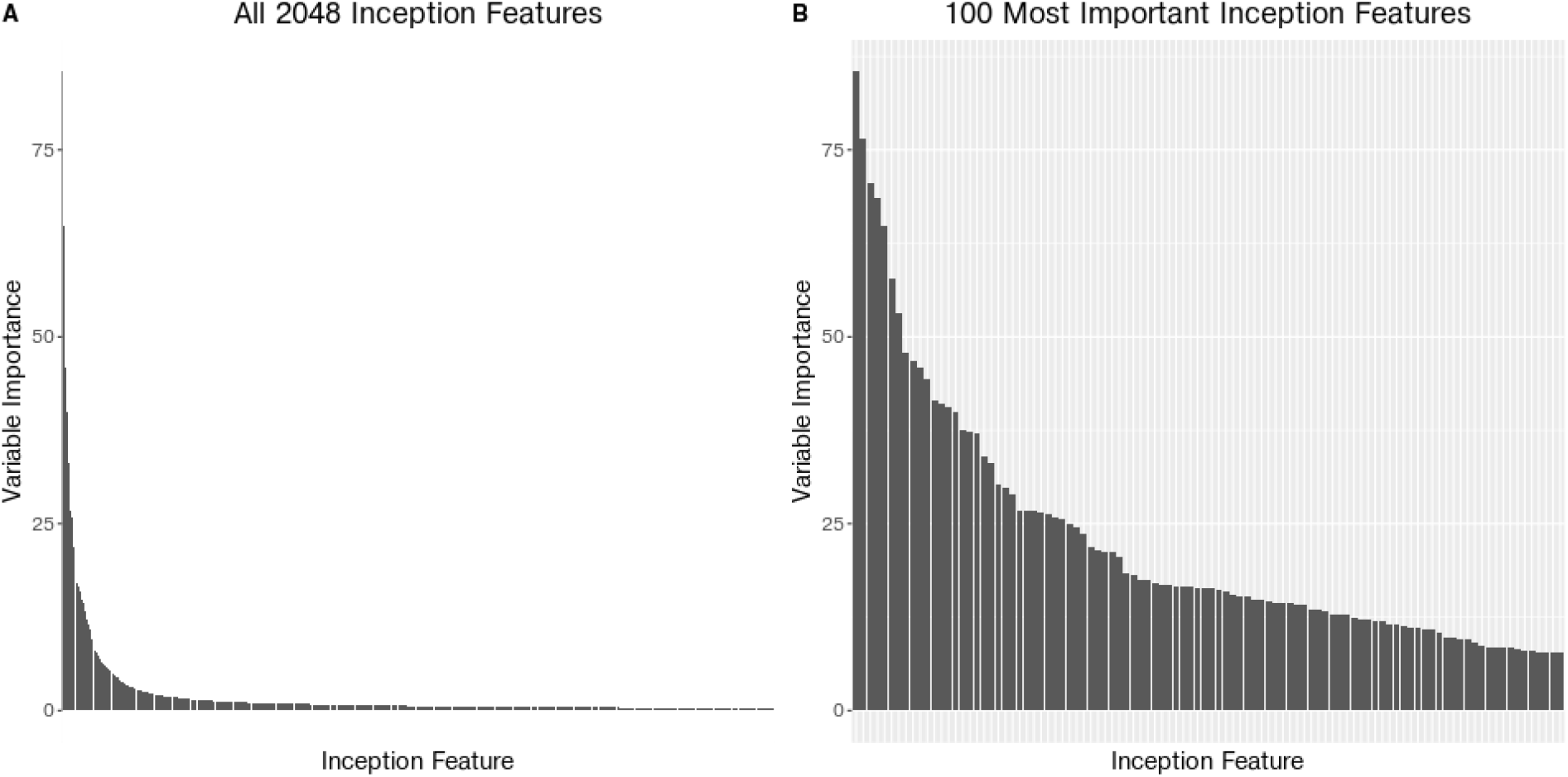
A small proportion of Inception features are important in classifying XTLD. Variable importance of Inception-based random forest classifier for (A) all Inception features and (B) the top 100 Inception features.

## Discussion

We presented a repository of nearly 1,000 H&E images of PDX samples and their human progenitors. All images are clinically annotated, many have paired expression and genomic data, and several have associated tumor, stromal, necrosis regions manually segmented by an expert pathologist. To the best of our knowledge, this is the first, large-scale, publicly-accessible set of PDX-derived images. As a demonstration of the utility of this resource, we provided two use cases: (1) a tumor-associated region classifier and (2) a classifier of XTLD – the unintended outgrowth of human lymphocytes, rather than the epithelial tumor cells, at the xenograft implantation site.

The images and all associated transcriptomic and genomic data, metadata, and annotations are hosted on the Seven Bridges Cancer Genomics Cloud [CGC; www.cancergenomicscloud.org; (Lau et al. 2017)]. The CGC colocalizes data and computational resources, including those with graphics processing units (GPUs), in the cloud to enable efficient analysis. We have implemented the major components of our processing pipeline, namely quality control and tissue mask generation with HistoQC and cell segmentation and calling with Hover-Net, as publicly-available “apps” – effectively containers that can be automatically distributed across computational cloud resources for parallel processing of images. All analysis and processing for this manuscript was performed on the CGC. Practical execution of Hover-Net benefited from the CGC, as it is computationally intensive and requires GPUs. By the time of publication, the images will also be hosted on the Imaging Data Commons (Fedorov et al. 2021) and all genomic and transcriptomic data are hosted on the PDXNet portal (Koc et al. 2022).

The repository enables exploration of several key biological questions, many specifically relevant to PDXs: (1) What morphological features of human tumors are recapitulated in PDXs? (2) Stromal cells and cancer-associated fibroblasts (CAFs), in particular, have been implicated in disease largely through their (highly varied) mediation of tumor / immune cell interactions (Mao et al. 2021). Do they retain this or some other function in immunocompromised mice? How are answers to these impacted by passaging, with early passages still harboring human immune and stromal cells and with murine CAFs replacing their human counterpart over time and passages (Blomme et al. 2018). Addressing these would be facilitated by placing immune and stromal cells in the context of the tumor core, periphery, or exterior using our regional classifier. (3) Automated grading of prostate (Nir et al. 2019), breast (Ehteshami Bejnordi et al. 2018), and colon (Vuong et al. 2020) cancers can be achieved through deep learning-based analysis of H&E images. Is tumor severity similarly imprinted on PDX sample morphology? If so, it should be possible to use the H&E images in our repository to predict their associated tumor stage and/or grade annotations. (4) Chemotherapy-treated breast samples exhibit morphological differences relative to treatment naive samples, including enlarged tumor cell nuclei (Rasbridge et al. 1994). Are such differences observed across tumor types and in PDX models? (5) Genetic mutations (Coudray et al. 2018) and gene expression (Schmauch et al. 2020) have been predicted from H&E using deep learning approaches. The (bulk) whole-exome sequencing and RNA-seq data paired with histology images in our repository would provide labels for transcriptome-wide studies in PDX samples. What gene mutations and expression patterns can be predicted? Do they validate published results (Coudray et al. 2018; Schmauch et al. 2020)? Are the associated, predictive morphological signatures preserved across tumor types?

Our region and XTLD classifiers were trained on features engineering from cells segmented and classified by one particular algorithm – Hover-Net. As a demonstration of the general applicability of our annotated images, we applied a second approach, Histology-based Digital (HD)-Staining, to a lung sample (S. Wang et al. 2020). HD-Staining has proven effective in classifying a rich set of cell types, including a tripartite separation of immune cells into macrophages, red blood cells, and a broader immune category. Intriguingly, we observed significant differences between Hover-Net- and HD-Staining-predicted cell types within stromal and necrotic regions annotated by our pathologist. These region-level annotations make possible a more systematic comparison of Hover-Net, HD-Staining, and related methods across other tumor types (Shmatko et al. 2022). This remains for future work.

H&E image color variability observed across institutions and labs (Nam et al. 2020; Otálora et al. 2019; Jose et al. 2021) may be caused by differences between stain batches and manufacturers, staining and fixation protocols, and tissue thickness (Veta et al. 2014; Bancroft 2008, chap. 10), as well as differences across imaging parameters, including scanner models (Leo et al. 2016) and image magnification (Komura and Ishikawa 2018). These site-specific effects adversely impact downstream DL-based classification (Shafiei et al. 2020; Salvi, Michielli, and Molinari 2020; Zheng et al. 2019) and confound prediction of survival, mutations, and tumor stage (Howard et al. 2021). The resulting lack of generalizability is a significant barrier to clinical deployment of these approaches (van der Laak, Litjens, and Ciompi 2021). Several classes of color normalization attempt to address this issue: (1) template color matching maps summary statistics or histograms of RGB values in a reference image to those in a reference image (Reinhard et al. 2001; Kothari et al. 2011; Y.-Y. Wang et al. 2007; Magee et al. 2009; Janowczyk, Basavanhally, and Madabhushi 2017); (2) color deconvolution (Ruifrok and Johnston 2001) represents each of the H&E dyes as a “stain vector” and substitutes a reference stain vector for the corresponding target vector (Macenko et al. 2009; Rabinovich, A. and Agarwal, S. and Laris, C. A. and Price, J. H. and Belongie, S. n.d.; Trahearn et al. 2015; Magee et al. 2009; Khan et al. 2014; Shafiei et al. 2020; Salvi, Michielli, and Molinari 2020; Zheng et al. 2019; Vahadane et al. 2016); and (3) generative adversarial network (GAN) approaches transfer strain distributions from a reference dataset (rather than single image) to a target dataset (Bentaieb and Hamarneh 2018; Tarek Shaban et al. 2018; Wagner et al. 2021). Our repository has considerable variability across the dimensions impacting image color, with six sites contributing data generated at two resolutions (20X and 40X) from different scanner models across multiple years. As such, it offers ample opportunity to compare color normalization approaches.

The H&E images presented here, alongside their genomic, transcriptomic, clinical, and pathological data, complement existing large-scale repositories, including TCGA, TCIA (Prior et al. 2013), GTEx (GTEx Consortium 2013), and CAMELYON17 (Litjens et al. 2018; Ehteshami Bejnordi et al. 2017; Bandi et al. 2019), while uniquely contributing PDX samples. As such, they expand the heterogeneity of publicly-accessible histology data available for technical studies (e.g., color normalization and batch correction), while also enabling exploration of the phenotypic effect of cellular interactions and morphology within a model system specifically intended to capture them.

## Methods

### Mapping imaging and genomic data

Images and genomic data are identified at the patient, model, and sample level. These levels are mapped using the following columns in the imaging and sequencing metadata for PDMR:

Imaging and sequencing mapping for PDMR data

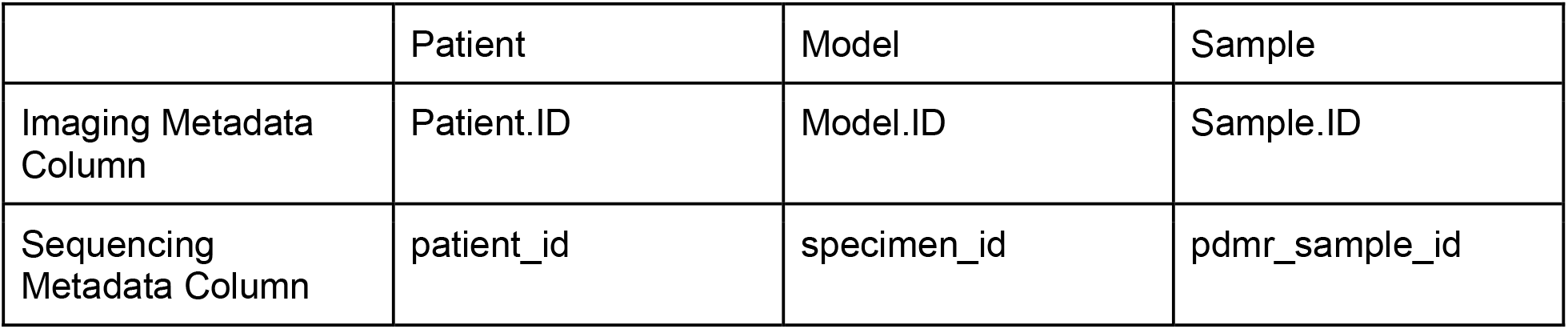

|  | Patient | Model | Sample |
| --- | --- | --- | --- |
| Imaging Metadata Column | Patient.ID | Model.ID | Sample.ID |
| Sequencing Metadata Column | patient_id | specimen_id | pdmr_sample_id |

Levels are mapped in the PDTCs using the following translations:

Imaging and sequencing mapping for PDTC data

|  | Patient | Model | Sample |
| --- | --- | --- | --- |
| Imaging Metadata Column | Patient.ID | Model.ID | Sample.ID |
| Sequencing Metadata Column | patient_id | model_id | sample_id |

### Image processing

For H&E images analyzed with Hover-Net or Inception, quality control and tissue segmentation were first performed using HistoQC (Janowczyk et al. 2019). Cell nuclei were then segmented and labeled as neoplastic, necrotic, immune, stromal, or other using Hover-Net (Graham et al. 2019). Features of 512 × 512 pixel tiles at 20X resolution (0.50 um per pixel) were computed using the Inception V3 convolutional neural network pre-trained on ImageNet (“Rethinking the Inception Architecture for Computer Vision” n.d.), as previously described (Noorbakhsh et al. 2020).

For H&E images analyzed with HD-Staining, tissue segmentation was first performed using Otsu thresholding followed with morphological dilation and erosions (Wang et al, 2018 https://www.nature.com/articles/s41598-018-27707-4). For whole slide analysis at 20X resolution, a 256 × 256 pixel window was slid over the entire tissue area with step size 226 pixel. Cell nuclei were then simultaneously segmented and classified as tumor, stroma, lymphocyte, red blood cell, macrophage, and karyorrhexis using HD-Staining (Wang et al, 2020). Only the nuclei with centroids located within the central 226 × 226 pixel area were kept to minimize the edge effect.

### Predictive analyses

Regional and XTLD random forest classifiers were trained and evaluated using five-fold cross validation with the ranger R package (Wright and Ziegler 2017).

